# Two-step voltage-sensor activation of the human K_V_7.4 channel and effect of a deafness-associated mutation

**DOI:** 10.1101/2025.02.04.636443

**Authors:** Mario Nappi, Damon J. A. Frampton, Ali S. Kusay, Kaiqian Wang, Francesco Miceli, Sara I. Liin, Maurizio Taglialatela, Antonios Pantazis

**Author notes:** Correspondence to: Maurizio Taglialatela & Antonios Pantazis **Email:**.

## Abstract

Voltage-gated, potassium-selective K_V_7.4 channels are expressed in the inner ear and are crucial for hair-cell function and survival. Loss-of-function variants of *KCNQ4*, the gene encoding K_V_7.4-channel subunits, cause non-syndromic progressive hearing loss (DFNA2). K_V_7.4 opening requires a voltage-dependent conformational change (activation) of the charged voltage-sensor domains (VSDs), and its transduction to the pore. Previously, fast charge displacement was reported during VSD activation at negative potentials, but it is unclear how this is coupled to slow channel opening occurring at more depolarized potentials. Here, we optically tracked K_V_7.4 VSD activation with voltage-clamp fluorometry, leveraging two different fluorophores and pulsed excitation, to thoroughly characterize VSD movements. We found that VSD activation comprises several voltage-dependent transitions, some of which had kinetics and voltage-dependence matching those of channel opening and closing. The deafness-associated mutation R216H, which substitutes a charged amino-acid in the VSD, impaired both VSD movements and channel opening, shifting them towards more depolarized potentials. This suggested that R216H impaired K_V_7.4 function by destabilizing VSD activation. Using molecular dynamics, we found that H216 reduced intramolecular interactions, thus decreasing the stability of an active VSD conformation. We propose that the K_V_7.4 VSD activates in two steps: a fast movement at negative voltages that represents a first transition to an intermediate state of activation; and this is followed by slower, depolarized component that represents subsequent full VSD activation, which drives channel opening.

## Introduction

*KCNQ1-KCNQ5* genes encode five distinct subunits of voltage-gated potassium channels, designated K_V_7.1 through K_V_7.5. Among them, K_V_7.4-channel subunits are expressed in inner and outer hair cells (IHCs and OHCs) of the cochlea and in central auditory pathways of the brainstem (1, 2), as well as in skeletal muscle (3) and vascular smooth muscle cells (4). Loss-of-function (LoF) mutations in the *KCNQ4* gene cause non-syndromic autosomal dominant deafness (DFNA2) (1), a progressive form of sensorineural hearing loss.

Each K_V_7.4 subunit is comprised of six transmembrane segments (S1-S6), with the voltage-sensor domain (VSD) formed by S1-S4, while the pore domain is comprised of S5 and S6 (Fig.1A). These subunits assemble into functional channels, either as homotetrameric or heterotetrameric complexes with other K_V_7-family members (5). Channel opening occurs in response to membrane depolarization, whereupon the positively charged S4 helix (Fig.1B) moves outward to its activated state. This conformational change is then transmitted to the pore, a process referred to as “electromechanical coupling”, opening the gate and permitting K^+^ flow (6).

**Figure 1:**
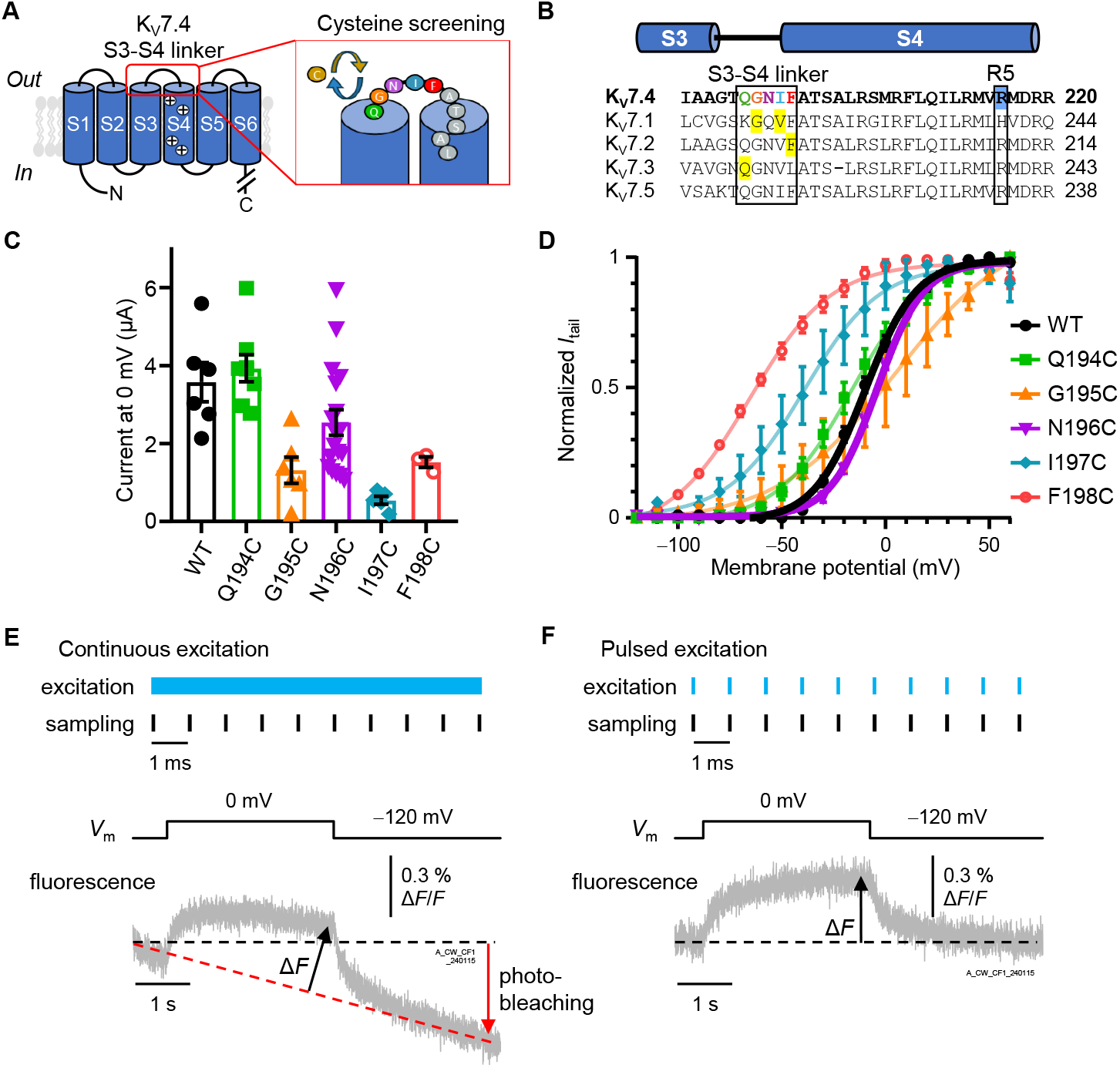
Setting the stage for VCF experiments on K_V_7.4. **(A)** K_V_7.4-subunit membrane topology and close-up on the S3-S4 linker. Four subunits assemble to form a functional channel. VSD = voltage-sensor domain; PD = pore domain. Cysteine scanning mutagenesis was performed for every residue in the linker. **(B)** Amino-acid sequence alignment for all human K_V_7 channels, showing the S3-S4 linker and adjacent regions. Amino-acid positions labelled for VCF in studies of other K_V_7 channels (7, 8, 11, 13) are highlighted in yellow. R216 is highlighted in blue. Homologous amino acids are indicated in the black box. **(C)** Comparison of current amplitudes for K_V_7.4 WT and S3-S4-linker mutants. **(D)** Mean, normalized voltage-dependence of channel opening in K_V_7.4: WT (grey); Q194C (green); G195C (orange); N196C (purple); I197C (light blue); F198C (red). *n* = 3-18 cells. Voltage-dependence parameters and statistical analysis in Table S1. The N196C substitution had minimal functional effects and was chosen for further analysis. It is thenceforth referred to as K_V_7.4*. **(E)** In “traditional” VCF protocols, fluorescence excitation is constantly applied (blue bar). Over time this causes photobleaching, i.e. light-induced fluorophore damage. Between data sampling (every 1 ms), no emission is recorded, but photobleaching still occurred. Grey trace shows fluorescence from K_V_7.4* labelled with AF488. The voltage pulse resulted in a fluorescence change (Δ*F*, reporting VSD activation). The fluorescence decayed due to photobleaching (red dashed line). **(F)** In a pulsed VCF protocol, the excitation LED was on 10% of the total time (100 μs every 1 ms), resulting in 90% less bleaching, as compared to the fluorescence trace from the same cell as in (E). Error bars are SEM.

VSD activation has been characterized in some members of the K_V_7 family (7–13). The K_V_7.1 VSD activates in two steps, first transitioning from resting to intermediate and then to the fully activated state. Both the intermediate and fully-activated states are coupled to distinct conducting states, with different biophysical and pharmacological properties (14, 15, 10, 12). In contrast to K_V_7.1, the VSD of K_V_7.2 and K_V_7.3 channels does not assume an intermediate state (13, 16). In several previous studies, disease-associated mutations in K_V_7 channels have been utilized to inform on the gating mechanism of the channels and provide a molecular etiology (11, 13, 16–18). Using gating-current measurements, we previously reported that the K_V_7.4 VSD activates faster and at potentials substantially more negative compared to channel opening (19). These results raised a conundrum, of how fast and “left-shifted” VSD movements relate to the slow and depolarized K_V_7.4-channel opening. Given the subtype-specific diversity in voltage-dependent gating among the K_V_7 channels studied thus far, inferring VSD operation in K_V_7.4 based on the activation mechanisms in K_V_7.1-K_V_7.3 is not possible. To better characterize K_V_7.4 VSD activation and its coupling to channel opening, we used voltage-clamp fluorometry (VCF) to optically track VSD movements in conducting K_V_7.4 channels. We performed VCF experiments in the absence or presence of R216H: a LoF DFNA2-associated mutation (20) that removes a putative voltage-sensing charge in S4. Finally, we complemented our studies with structural simulations.

## Results

### Implementing VCF on K_V_7.4 channels

In most VCF studies of voltage-gated ion channels, an environment-sensitive, thiol-reactive fluorophore is conjugated to a substituted cysteine at the S3-S4 extracellular linker. When the charged S4 traverses the membrane electric field upon depolarization, the environment of the attached fluorophore will change, and so will its fluorescence emission. In this way, VSD activation can be tracked as ensemble fluorescence deflections (Δ*F*) (21– 26). To select a labelling site appropriate for VCF recordings in K_V_7.4, we engineered five cysteine mutants in the S3-S4 loop (Q194C, G195C, N196C, I197C and F198C, Fig.1A,B) and used two-electrode voltage clamp to compare their function to that of the wild-type channel. We selected N196C for further study as it did not reduce current levels and only induced a modest shift in voltage dependence of approximately +10 mV (Fig.1C-D, Table S1). K_V_7.4(N196C) channels are henceforth referred-to as K_V_7.4*.

Labelling K_V_7.4* with Alexa Fluor 488 C_5_ Maleimide (AF488) revealed robust Δ*F* (Fig.1E). However, the prolonged excitation resulted in strong fluorescence photobleaching, which was challenging to effectively subtract for analysis. To limit the impact of photobleaching on the fluorescence signal we developed a pulsed excitation protocol for VCF. This maneuver eliminated most photobleaching, while leaving Δ*F* amplitude intact (Fig.1F).

No Δ*F* were detectable in WT K_V_7.4 labelled with AF488, confirming that the Δ*F* arose from fluorophores attached to C196, just outside S4 (Fig.S1A). WT K_V_7.4 labelled with AF488 exhibited a slightly hyperpolarized voltage dependence compared to unlabelled WT channels (Fig.S1). Combined with the slightly depolarized opening of K_V_7.4* (Figs.1D, S1), we later show that the voltage dependence of labelled K_V_7.4* channel opening was practically wild-type-like.

### The K_V_7.4 VSD activates during closed states and exhibits two kinetic components

Having identified an ideal labelling position and VCF protocol, we proceeded to characterize the VSD movements, as reported from Δ*F* signals, and relate them to channel opening (Fig.2). Voltage dependence was determined by fitting Δ*F* (VSD activation, or “F(V)”) and tail current (channel opening, or “G(V)”) to the Boltzmann distribution (Fig.2C). Interestingly, VSD activation had an apparent voltage-dependence shifted by ∼−20 mV relative to that of channel opening (Fig.2C). A straightforward interpretation of this result is that the VSD undergoes conformational changes during closed states of the channel.

**Figure 2:**
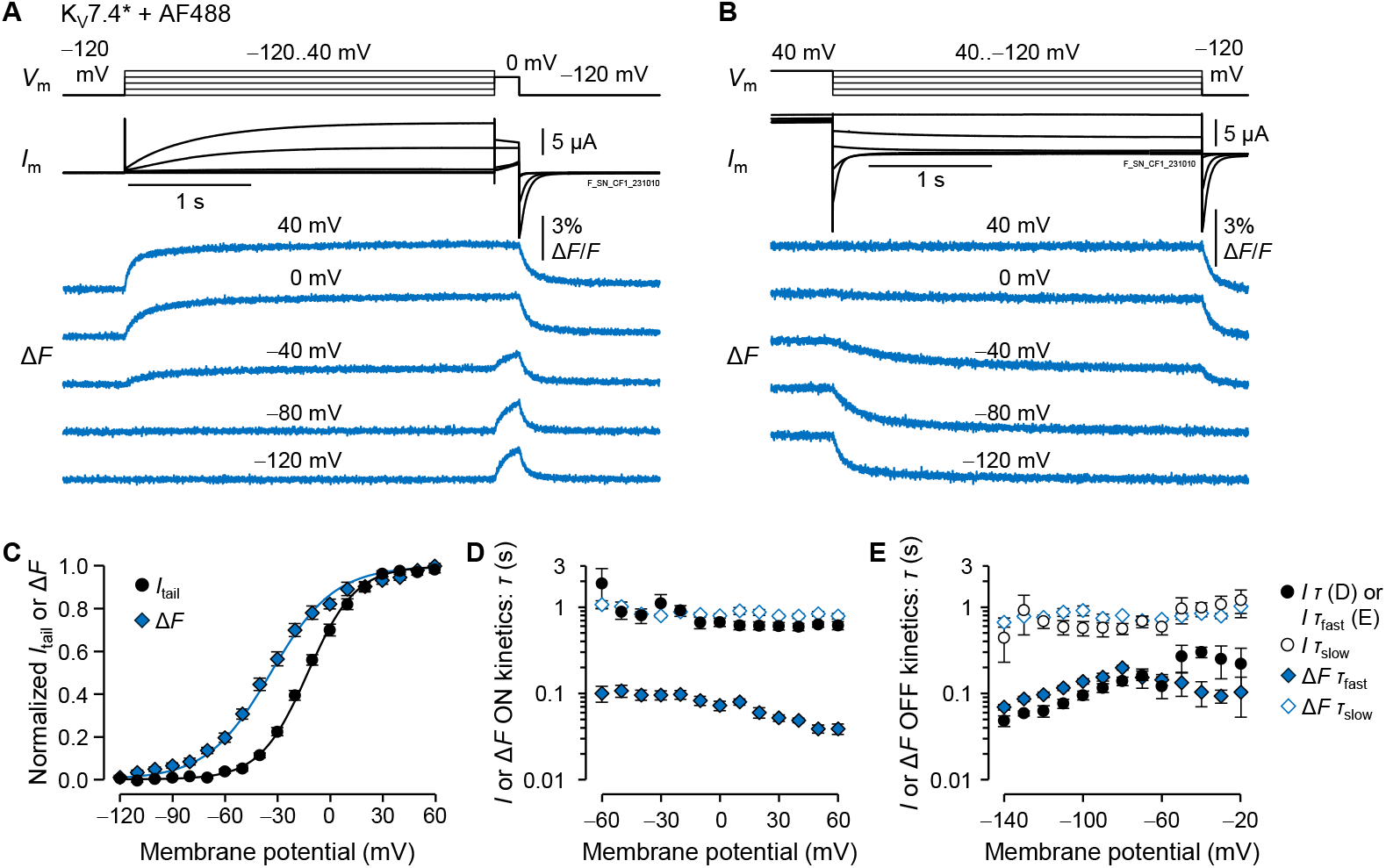
The K_V_7.4 VSD activates at negative potentials compared to channel opening. **(A)** Traces from an exemplary experiment showing the voltage protocol (*V*_m_), current (*I*_m_) and simultaneously acquired fluorescence signals (Δ*F*) from K_V_7.4* channels labeled with AF488. **(B)** As in (A) from the same cell, with a protocol to study channel closing and VSD deactivation. **(C)** Mean, normalized voltage-dependence of channel opening (*I*_tail_, filled black circles) and VSD activation (Δ*F*, filled blue diamonds) in K_V_7.4* labelled with AF488. Curves represent fits to Boltzmann distributions (eq.2,3). *I*_tail_: *V*_0.5_ = −13 ± 1.6 mV; *z* = 1.9 ± 0.085 *e*_0_; *n* = 11 cells; Δ*F*: *V*_0.5_ = −34 ± 2.1 mV; *z* = 1.4 ± 0.11 *e*_0_; *n* = 15 cells. The Δ*F* signals developed earlier in voltage compared to channel opening. **(D)** Time constants (*τ*) of channel opening (filled black circles) and VSD activation. Note that the latter had two kinetic components: a fast (*τ*_fast_, filled blue diamonds) and a slow one (*τ*_slow_, open blue diamonds); *τ*_slow_ was similar to the rates of channel opening. **(E)** Time constants (*τ*) of channel closing (fast: filled black circles; slow: open black circles) and VSD deactivation (fast: filled blue diamonds; slow: open blue diamonds). The fractional amplitudes of the kinetic components are in Fig.S2A,B. Error bars are SEM.

Kinetics were determined by fitting current or Δ*F* to single, or the sum of two, exponential distributions. VSD activation exhibited two kinetic components, one fast (*τ*_fast_∼1-10 ms) and one slow (*τ*_fslow_∼1 s; Figs.2D & S2A). Channel opening was dominated by one component, which overlapped with the slow fluorescence component (Fig.2D). Both VSD deactivation and channel closing exhibited two kinetic components which overlapped with each-other (Fig.2E & S2B).

### The K_V_7.4 VSD activates in two steps with distinct voltage dependence

The presence of two kinetic components for VSD activation and deactivation indicated that the voltage sensor undergoes distinct voltage-dependent transitions; however, this was not obvious in the apparent voltage-dependence of the AF488 Δ*F* signal (Fig.2C). We hypothesized that an alternative fluorophore may report a composite VSD voltage-dependence more clearly. Accordingly, we repeated experiments in K_V_7.4* using the 6-TAMRA C6 maleimide (TAMRA C6) fluorophore (Fig.3). In addition to slightly different photochemistry (its excitation and emission wavelengths are red-shifted compared to AF488), the TAMRA C6 fluorophore possesses an additional carbon link between the fluorescent and maleimide (cysteine-binding) moieties, presumably affording it greater length and flexibility (27).

TAMRA C6 Δ*F* (Fig.3A) were qualitatively different from those of AF488 (Fig.2A). Whereas AF488 Δ*F* increased monotonically with progressive depolarization, TAMRA C6 produced a more complex signal: modest depolarizations (up to ∼−30 mV) resulted in fluorescence decrease (Δ*F* < 0); in stronger depolarizations, an additional, slower fluorescence component became apparent with opposite amplitude to the first (Δ*F* > 0). (Fig 3A,B). WT K_V_7.4 showed no Δ*F* when labelled with TAMRA C6 (Fig.S3), confirming that the composite Δ*F* arose from labelling C196. Intriguingly, the two TAMRA C6 fluorescence components had similar kinetics to those of AF488; and the slower component had similar kinetics to channel opening (Figs.3C & S2C).

**Figure 3:**
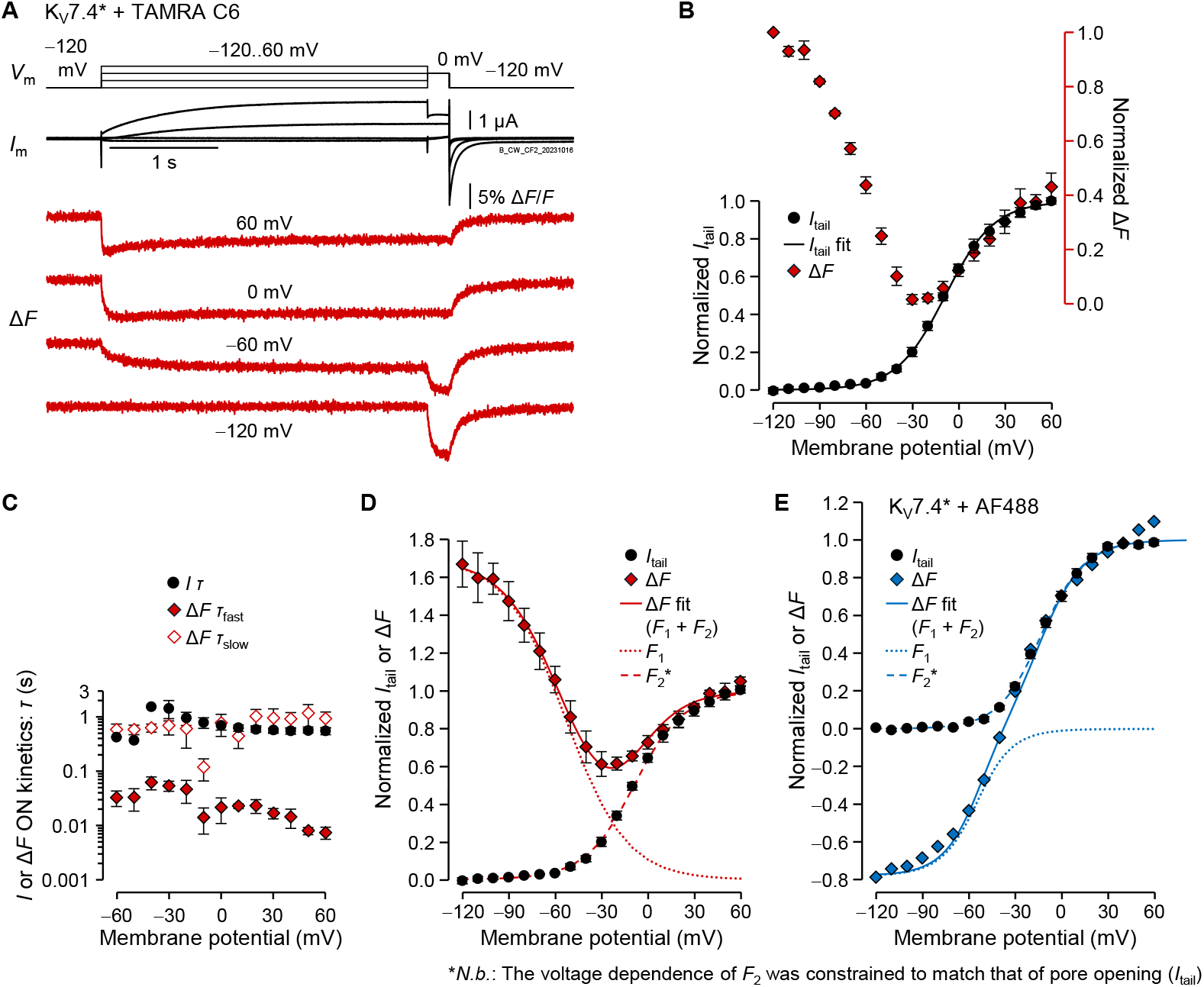
A second VSD transition is resolved at voltages relevant to channel opening. **(A)** The voltage protocol (*V*_m_, black), along with current (*I*_m_, black) and fluorescence (Δ*F*, red) traces from an exemplary experiment with K_V_7.4* labelled with TAMRA C6. **(B)** Mean, normalized channel opening voltage-dependence for K_V_7.4* labelled with TAMRA C6 (*I*_tail_, black filled circles). The black curve is the fit to a Boltzmann distribution (eq.2): *V*_0.5_ = −8.6 ± 0.89 mV; *z* = 1.6 ± 0.32 *e*_0_; *n* = 3 cells. Mean Δ*F* amplitude (normalized to its minimal and maximal values; red filled diamonds; plotted on the red scale; *n* = 3) showed two fluorescence components. The Δ*F* plot has been scaled and positioned so the second component aligns with the current voltage-dependence. **(C)** Time constants (*τ*) of channel opening (filled black circles) and VSD activation. Note that the latter had two kinetic components: a fast one (*τ*_fast_, filled red diamonds) and a slow one (*τ*_slow_, open red diamonds); the latter was similar to channel opening. The fractional amplitudes of the kinetic components are in Fig.S2C. **(D)** Mean Δ*F* (red filled diamonds) fit to the sum of two Boltzmann distributions (eq.3; solid red curve). *F*_1_ (dotted red curve): *V*_0.5_ = −53 ± 2.9 mV; *z* = 1.3 ± 0.029 *e*_0_; 63 ± 1.8% fractional amplitude. *F*_2_ (dashed red curve) was constrained to have the same voltage dependence as channel opening (panel B). The Δ*F* data are normalized to the *F*_2_ component, and the *I*_tail_ data are shown super-imposed (filled black circles). **(E)** AF488 Δ*F* (filled blue diamonds; from Fig.2) could also be expressed as the sum of two voltage-dependent components (eq.3; solid blue curve). *F*_1_ (dotted blue line) developed at negative potentials, as with TAMRA C6: *V*_0.5_ = −55 ± 1.3 mV; *z* = 2.3 ± 0.16 *e*_0_; 42 ± 3.8% fractional amplitude; *n* = 11; *F*_2_ (dashed blue curve) was constrained to have the same voltage dependence as channel opening (Fig.2C). As in (D), the Δ*F* data were normalized to *F*_2_ and the *I*_tail_ data from AF488-labeled K_V_7.4* channels are superimposed (filled black circles). Error bars are SEM.

We surmised that, since the slow Δ*F* component had similar kinetics to channel opening, it should also have similar voltage dependence. Accordingly, we fit the voltage dependence of TAMRA C6 Δ*F* to the sum of two Boltzmann distributions. The parameters of the first (*F*_1_) component were free, while the second (*F*_2_) component was constrained to have the same voltage-dependence parameters (half-activation potential, *V*_0.5_, and valence, *z*) as channel opening in the same cell. This resulted in an excellent fit to the data (Fig.3D), with *F*_1_ having a very negative voltage dependence (*V*_0.5_ = −53 mV). The two components had similar valence and *F*_1_ accounted for 60% of the total Δ*F*.

Thus, use of the TAMRA C6 fluorophore more clearly showed that the K_V_7.4 VSD activates in two sequential steps, each with distinct voltage-dependence and kinetics. If the two fluorophores tracked the same VSD movements, why did the voltage-dependence of AF488 exhibit a single component? We hypothesized that AF488 also reported two voltage-dependent components, but they were difficult to discern. This hypothesis was supported by a re-evaluation of the AF488 voltage-dependence with the sum of two Boltzmann distributions, one of them possessing the same voltage-dependence as channel opening (Fig.3E). The two activation transitions of the K_V_7.4 VSD were reported by both fluorophores, albeit differently.

### R216H impairs both VSD activation transitions and channel opening

How relevant are the K_V_7.4 VSD transitions to channel opening? We sought to characterize the impact of a deafness-associated mutation, R216H (20), on VSD activation and channel opening, for two reasons: first, to provide a molecular etiology for the disease, and second, because the variant produces an amino-acid substitution at a VSD position likely important for its activation. R216 is a conserved, positively charged arginine in S4, homologous to the fifth arginine at the S4 of the archetypal Shaker voltage-gated K^+^ channel (“R5”, Fig.1B). Thus, this clinically relevant variant could be used as a biophysical tool, to probe the link between VSD activation and channel opening.

VCF data from K_V_7.4* R216H channels (Fig.4A-C) were qualitatively similar to those from K_V_7.4* (Figs.2A-B, 3A). Our analysis revealed that R216H impaired the voltage dependence of channel opening and both VSD activation components (Fig.4D-F). Specifically, channel opening and *F*_2_ (which was constrained to have the same voltage-dependence) were shifted by 17 or 26 mV to more positive potentials, depending on whether the AF488 or TAMRA C6 label was used. *F*_1_ was also affected, showing 15-20 mV shifts. R216H also reduced the effective valence of these processes, by 10-25%. Consistent with this analysis, we also report a shift by ∼10 mV when AF488 Δ*F* data were fit to a single Boltzmann curve, concomitant with a 10% valence reduction (Fig.S2D). A straightforward interpretation of this analysis is that R216H hampered both VSD activation components and channel opening, such that they required more depolarization to occur (right-shifted *V*_0.5_); and reduced their sensitivity to changes in voltage (reduction in effective valence).

**Figure 4:**
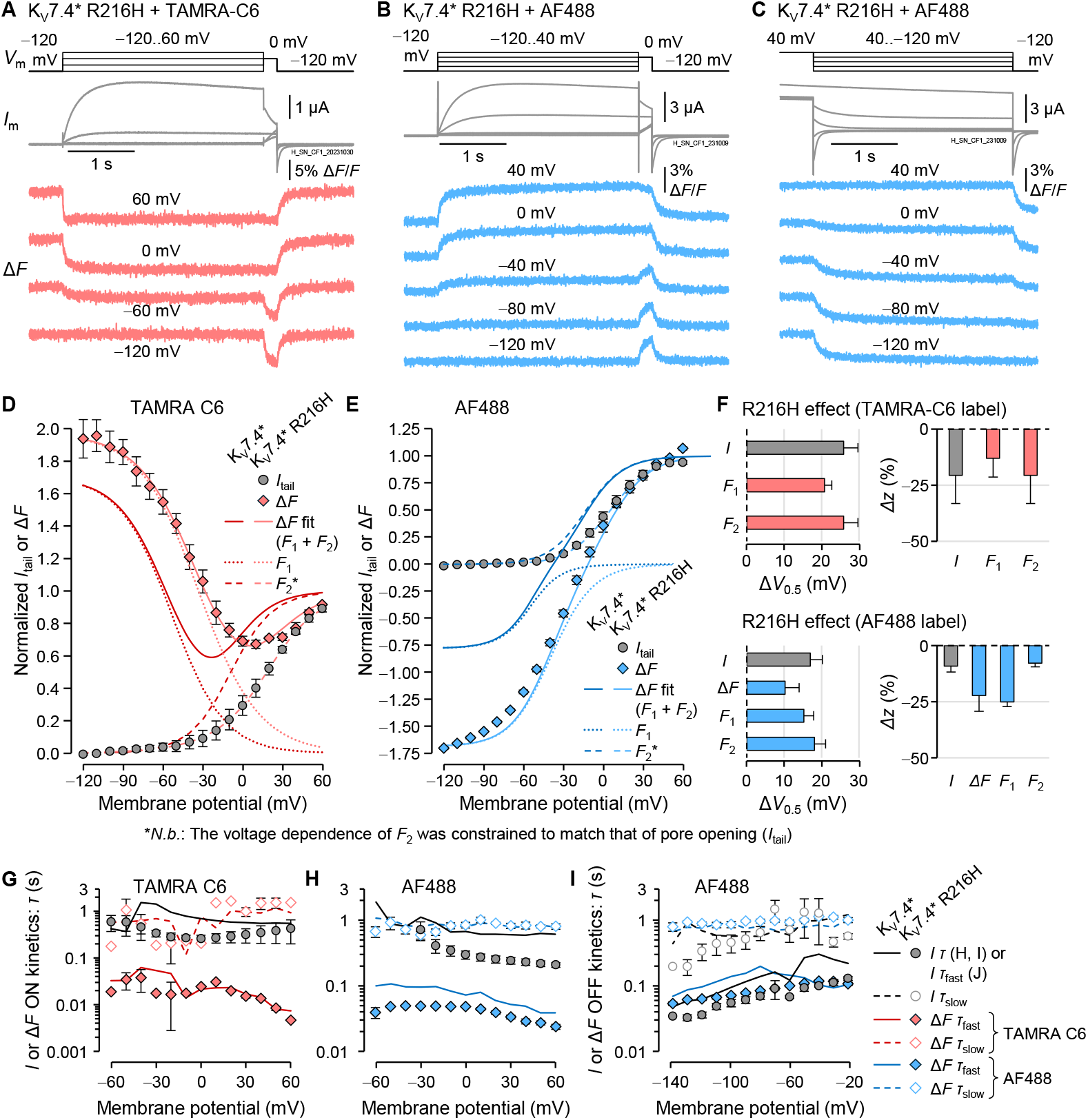
R216H impairs both VSD activation transitions and channel opening. **(A)** Traces from an exemplary experiment showing the voltage protocol (*V*_m_), current (*I*_m_) and simultaneously acquired fluorescence signals (Δ*F*) from K_V_7.4* R216H channels labeled with TAMRA C6. **(B)** As in (A), showing traces from a cell K_V_7.4* R216H channels labeled with AF488. **(C)** Traces from the cell in (B), with a protocol to study channel closing and VSD deactivation. **(D)** Mean, normalized voltage-dependence of channel opening (*I*_tail_, filled grey circles) and VSD activation (Δ*F*, filled light-red diamonds), from K_V_7.4* R216H channels labeled with TAMRA C6. The solid light-red curve represents a fit of the Δ*F* to the sum of two Boltzmann distributions (eq.3): *F*_1_ (dotted light-red line): *V*_0.5_ = −32 ± 2.0 mV; *z* = 1.1 ± 0.11 *e*_0_; 66 ± 1.3% fractional amplitude; *n* = 3; *F*_2_ (dashed light-red curve) was constrained to have the same voltage dependence as channel opening (*I*_tail_, filled grey circles): *V*_0.5_ = 17 ± 3.8 mV; *z* = 1.3 ± 0.20 *e*_0_; *n* = 3. Note that the Δ*F* was normalized to the amplitude of the *F*_2_ component. The TAMRA-C6 Δ*F* voltage-dependence from K_V_7.4* channels is shown for reference: total Δ*F*: solid red curve; *F*_1_: dotted red curve; *F*_2_: dashed red curve (from Fig.3D). Note that the effects of R216H on the voltage dependence of channel opening can be appreciated by comparing the *F*_2_ curves and, more explicitly, in Fig.S2D. **(E)** As in (D), for K_V_7.4* R216H channels labeled with AF488. As with K_V_7.4* channels (Fig.3E), the monotonic AF488 Δ*F* could be approximated by the sum of two components (*F*_1_, *F*_2_), with the latter having the same voltage-dependence as channel opening. **(F)** Bar graphs showing the effects of R216H on K_V_7.4* voltage-dependence parameters: *V*_0.5_ shifts (Δ*V*_0.5_) and percentile changes in *z* (Δ*z*), for channel opening (*I*, grey) and VSD activation (light red or blue, for TAMRA C6 or AF488, respectively). In the latter case, the effects on Δ*F* on fit with a single Boltzmann distribution are also shown (see Fig.S2D). **(G)** Time constants (*τ*) of channel opening (filled grey circles) and VSD activation (slow: filled light-red diamonds; fast: open light-red diamonds) for K_V_7.4* R216H channels labeled with TAMRA C6. Data from K_V_7.4* channels are shown as lines (from Fig.3C). **(H)** As in (G), for K_V_7.4* R216H channels labeled with AF488. Lines show K_V_7.4* data from Fig.2D. **(I)** The kinetics of pore closing and VSD deactivation K_V_7.4* for K_V_7.4* R216H channels labeled with AF488. Lines show K_V_7.4* data from Fig.2E. The fractional amplitudes of the kinetic components are in Fig.S2E-G. Error bars are SEM.

We also compared the activation and deactivation kinetics of K_V_7.4* and the K_V_7.4* R216H mutant (Figs.4G-I, S2E-G). R216H apparently accelerated the kinetics of channel opening, although this measurement may have been affected by current inactivation, observed in this mutant (Fig.4A,B). We could not detect a consistent change in fluorescence kinetics for both fluorophores; however, the fast component from AF488 measurements was accelerated in both activation (Fig.4H) and deactivation, concomitant to channel closure (Fig.4I). It is possible that the opposite direction of the Δ*F* components observed by TAMRA C6 affected the kinetic measurements, especially when they were similar in amplitude.

### R216H decreases S4 positional stability

The voltage-dependence shifts induced by R216H suggest that the mutation decreases the stability of the active VSD states relative to the resting. To assign a structural interpretation to this result, we conducted molecular dynamics simulations of K_V_7.4 with the WT sequence and with the R216H mutant. Specifically, we used the apo K_V_7.4 structure (PDB: 7BYL) (28) with S4 in a depolarized conformation, likely corresponding an activated VSD state.

First, the structural stability of WT or R216H channels in our simulation framework was assessed using root mean square deviation (RMSD) of residues in the transmembrane segments. The RMSD plots indicate that within the first ∼ 200 ns of simulation, the proteins had an overall deviation of around 2.7 Å relative to the initial structures (Fig.S4), which is consistent with thermal noise. Subsequent analyses were conducted on simulation frames from 200 ns and above to avoid frames where the protein was still equilibrating.

To examine how R216H affected VSD structural stability, we measured the distances between the Cα of R216 or H216 and the Cα of F143 (likely the charge-transfer-center residue (29) of K_V_7.4) during our simulation (Fig.5A). There was a wider distribution of distances in the R216H mutant compared to WT, and both distributions were centered around 12.5 Å (Fig.5B). This indicates that the mutant position H216 fluctuated more relative to the charge transfer center than the WT R216. The same was true for the intracellular half of the S4 helix bearing either R216 or H216. We observed that RMSD was shifted further from the initial structures and had a wider distribution in the presence of the mutant (Fig.5C).

**Figure 5:**
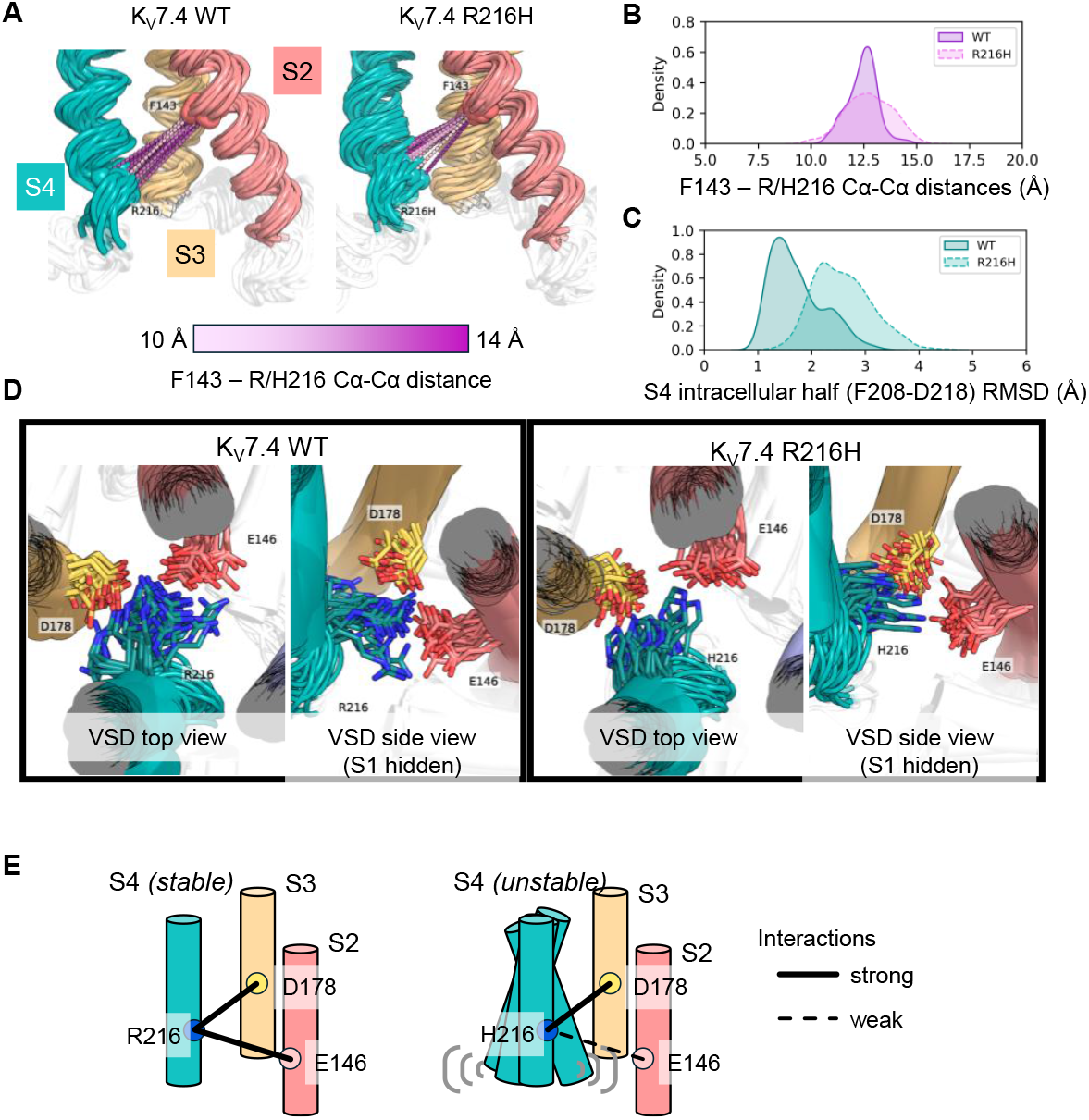
R216H destabilizes the intracellular part of S4. **(A)** Ensemble of simulation frames for K_V_7.4 with WT sequence or the R216H mutant. 10 frames were uniformly captured over the last 300 ns of three simulation replicas (30 frames in total). The Cα atoms of F143, R216 and R216H are illustrated as orbs. Distances between Cα atoms of F143 and R216 or H216 are shown as dashed lines with more intense purple shades representing longer distances. The S1 helix is hidden for clarity. **(B)** Distance distribution for F143 – R216 and F143 – H216, measured between the residue Cα atoms (top). **(C)** Root mean squared deviation (RMSD) distribution of Cα atoms in the S4 helix residues (F208-D218) measured after aligning on the S1-S3 helices (bottom). **(D)** Alternative views of simulations frame ensembles of K_V_7.4 to illustrate the sidechain conformations of E146, D178, and R216 or H216. The S1 helix is colored in light purple, the S2 helix in light red, the S3 helix in dark yellow and the S4 helix in teal. The simulations used the apo K_V_7.4 structure (PDB: 7BYL) (28). **(E)** Cartoon illustration of how the S4 helix is destabilized in the presence of H216 (right) compared the WT R216 (left) due to the weakening of its interaction with E146.

To rationalize the difference in S4 positional stability, we analyzed the propensities of R216 or H216 to form stabilizing, intramolecular interactions with other residues. In simulations with the mutant, H216 was assigned the doubly protonated, +1-charge tautomer, to enable salt-bridge interactions with neighboring acidic residues. R216 formed prominent salt-bridge interactions with E146 in S2 (80±11 % of simulation frames). By comparison, H216, which is also protonated, but shorter and more conformationally restricted than R216, interacted with E146 far less frequently (9±5 %). Conversely, both R216 and H216 interacted with D178 (S3) with similar frequency (76±11 and 67±10 % of simulation frames, respectively). Apart from E143 and D178, no other residues formed prominent interactions (>10 % of simulation frames) with R216 or H216. The lack of interaction of H216 with E146 meant that the mutation-bearing S4 was less stably anchored, indicating that the activated VSD state in the R216H variant is less stable than in WT. Notably, when H216 is in the neutral tautomeric form, all stabilizing salt-bridge interactions would be abolished.

## Discussion

In this work, we sought to shed light on the voltage-sensing mechanism of K_V_7.4 channels. This involved five crucial steps: 1) identification of N196, an ideal fluorescence labelling site on the S3-S4 extracellular linker, which produced minimal functional effects on channel voltage-dependent opening (Figs.1A-B); 2) pulsed excitation, which largely abolished photobleaching (Fig.1E-F); 3) use of two different fluorophores, which helped resolve distinct VSD activation transitions (Figs.2, 3); 4) use of the R216H deafness-associated mutation at an eminent part of the VSD as a tool, to probe the relevance of VSD transitions to channel opening (Fig.4) and (5) leveraging a K_V_7.4 structure to rationalize the effects of R216H with molecular dynamics simulations (Fig.5). Thus we optically tracked the conformational changes underpinning the voltage-dependent development of physiological K_V_7.4 currents, which contribute to, for instance, the repolarizing n-current in OHCs and determining the vascular tone (30–33).

### The voltage-sensing mechanism of K_V_7.4 channels

Our optical investigation revealed two movements of the K_V_7.4 VSD, with distinct kinetics and voltage dependence. It is not immediately clear how the two steady-state, voltage-dependence components (*F*_1,_ *F*_2_) correlate with the two kinetic components (*τ*_fast_, *τ*_slow_) observed during VSD activation and deactivation. Nevertheless, in channels labeled with TAMRA C6, both the depolarized (*F*_2_) and the slow (*τ*_slow_) components match the voltage-dependence and kinetics of channel opening, respectively (Fig.3C, D). We propose that *F*_1_ represents the first activation transition of the K_V_7.4 VSD, from a resting to an intermediate-active conformation; and *F*_2_ a second transition, from the intermediate-active to a full-active state, which is linked to channel opening. This is illustrated in a scheme in Fig.6A.

**Figure 6:**
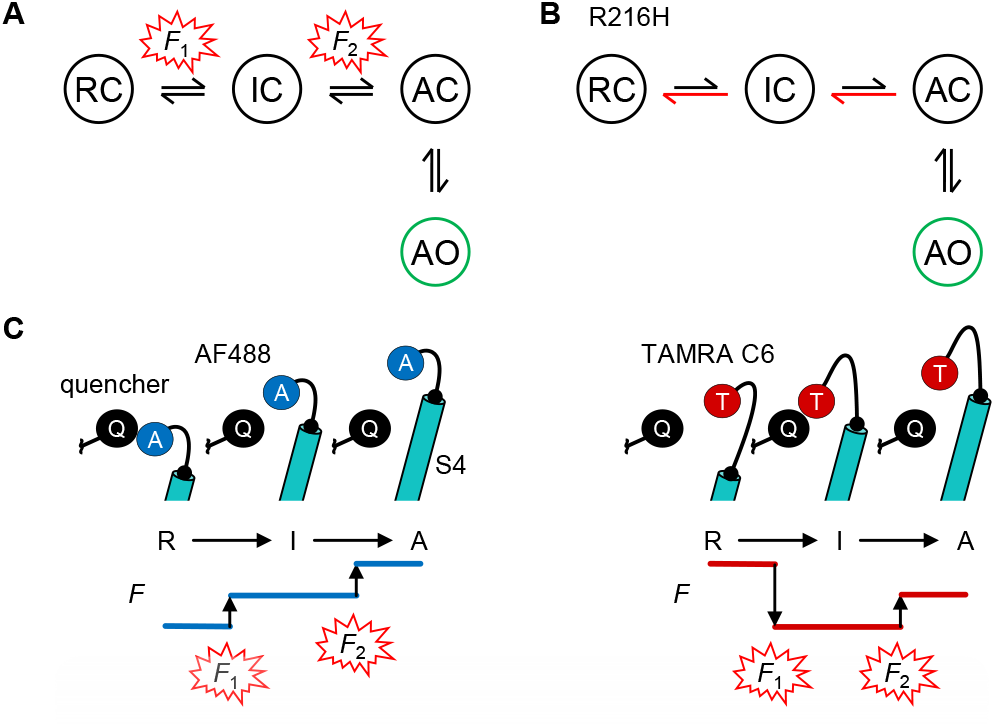
Proposed mechanisms of K_V_7.4 voltage-dependent activation, the effect of R216H, and VCF signals. **(A)** In this scheme of channel states, the VSD can assume three conformational states: resting (R), intermediate (I) and active (A). The pore can assume two possible conformational states: closed (C) or open (O). We propose that only the fully-active state of the voltage sensor induces opening of the pore. The R ↔ I and I ↔ A transitions have distinct voltage-dependence and give rise to the *F*_1_ and *F*_2_ components in the fluorescence signals. The vertical pore opening transition is thought to be voltage-independent **(B)** We propose that R216H destabilizes the intermediate- and fully-active VSD states, accelerating the VSD deactivation transitions (red arrows) and resulting in impaired VSD activation and channel opening. **(C)** Cartoon of a proposed mechanism for the differences in observed Δ*F* from AF488 and TAMRA C6 fluorophores. As S4 sequentially transitions from resting (R) to the intermediate (I) and active (A) states, it moves past a quenching group (Q). AF488 (left) is quenched in the R-state; as S4 activates, it moves farther from the quencher, which reduces its quenching probability and increases fluorescence emission. In this way, the macroscopic Δ*F* is monotonically increasing with voltage (Fig.2). TAMRA C6 (right) is more flexible, and thus experiences a different state-dependent interaction with the quencher, even while S4 undergoes the same movements as with AF488: the fluorophore is unquenched in the R-state; becomes more quenched in the I-state; and this quenching is partially relieved in the A-state. In this way, the macroscopic Δ*F* exhibits two macroscopic components with opposite amplitudes (Fig.3).

In this premise, the reason for relatively slow K_V_7.4 opening is the slow VSD activation, from the intermediate to the fully-active states (reported as *F*_2_). Another insight provided by studies of the R216H-mutant is the relevance of both VSD transitions to channel opening. The mutation results in the impairment (shift to more depolarized potentials) of both *F*_1_ and *F*_2_ VSD transitions, and channel opening (Fig.4D-F). MD simulations show that the mutation destabilizes an active VSD conformation, as H216 is less competent at forming stabilizing interactions with a counter-charge than R216 (Fig.5). This result is consistent with the accelerated VSD deactivation and pore closing observed in this mutant compared to wild-type (Fig.4I). Destabilization of active conformations (Fig.6B) accounts for the depolarizing shift in the steady-state voltage-dependence curves, and the loss-of-function phenotype, characteristic of deafness-associated variants of *KCNQ4*.

### Optical and electrophysiological methods to measure *K*_*V*_*7*.*4 gating*

Of note, the two VSD transitions were differentially reported by the two fluorescent probes. In the case of AF488, both VSD movements caused fluorescence unquenching (Figs.2A, 3E). In the case of TAMRA C6, *F*_1_ resulted in fluorescence quenching and *F*_2_ in unquenching (Fig.3A, D). It has been previously reported that different fluorophores may experience different state-dependent interactions with nearby quenchers and report different Δ*F* signals (27, 34–36). The two fluorophores here had subtle differences in photochemistry, length and flexibility: AF488 has a 5-carbon linker between the fluorescent and maleimide moieties; TAMRA C6 has a 6-carbon linker, so it is likely longer and more flexible, as previously shown for tetramethylrhodamine probes with variable linker lengths (27). Fig.6C shows a cartoon illustrating how the same S4 movements can be differentially reported by the two fluorophores, given the same quenching mechanism.

*F*_1_ is most similar to the charge displacement reported in a gating-current study in K_V_7.4 (19), as both develop faster and at more negative potentials than channel opening. *F*_2_ would have been extremely challenging to detect with gating currents: First, because the amplitude of gating currents is proportional to the rate of change movement, and *F*_2_ develops with a rate three orders of magnitude lower than more typically resolved charge movements (seconds instead of milliseconds). Second, because charge integration over the time-scales necessary to capture *F*_2_ would make measurements very susceptible to electrophysiological artifacts (noise, membrane instability). Third because, as *F*_2_ exquisitely tracks with channel opening, the pore blockade necessary for gating current measurement may allosterically alter *F*_2_ properties (or blockers may directly perturb VSD movements). While VCF circumvents these shortcomings, *F*_2_ was still challenging to detect optically, requiring confirmation with two fluorophores and a new method to decrease fluorescence photobleaching over the particularly long recording times, explained below.

### Technical interlude: pulsed excitation to minimize photobleaching

Fluorescence photobleaching occurs when a fluorophore in the short-lived excited state is struck by additional photons. In this case, the fluorophore attains higher energy levels, which may result in chemical modifications that abolish its ability to fluoresce. In VCF, photobleaching is evident as a decay of the optical signal over time. Here, slow K_V_7.4 kinetics (*F*_2_) necessitated the use of long recordings (up to 7 s), at which point the photobleaching matched or exceeded Δ*F* amplitude (Fig.1E).

To reduce photobleaching, we drew inspiration from a pulsed-excitation application in cardiac local-field fluorescence microscopy. In this work, Escobar and colleagues used high-power, picosecond laser excitation pulses, briefer than the mean fluorescence lifetime, to maximally excite rhod-2 fluorophores while minimizing secondary excitations and photobleaching (37). In our experiments, the slow K_V_7.4 kinetics allowed for a relatively low sampling rate (1 kHz). While our LEDs could not be pulsed at rates relevant to the fluorescence lifetime, it was possible to turn them on around the sampling point and keep them off for 90 % of the sampling interval, while data were not collected. In this way the K_V_7.4-conjugated fluorophores received full excitation during sampling, producing robust Δ*F*; while photobleaching was minimized, as the total light exposure was reduced by 90 % (Fig.1E, F). This was straightforward to implement in a commonly used Digidata/Clampex acquisition system. Even more precise LED control can be achieved with specialized timers, as shown in a pulsed-excitation VCF study of P2X7 channels (38).

### The K_V_7.4 voltage-sensing mechanism in context

Considering other members of the K_V_7-channel family, the K_V_7.4 voltage-sensing mechanism recapitulates most closely that of K_V_7.1 channels complexed with regulatory KCNE1 subunits (39): the VSD undergoes two sequential activation transitions, and the second is thought to be coupled to opening. Similar to our findings for K_V_7.4, only the second voltage sensor movement overlaps in voltage with channel opening. Another similarity between K_V_7.4 and K_V_7.1/KCNE1 channels is the slow time-course of channel opening. In the absence of KCNE1, K_V_7.1 channels can also open from the intermediate-active VSD state, do so much faster, and display F(V) and G(V) curves that largely overlap in voltage (although with noticeable *F*_1_ and *F*_2_ components). Largely overlapping F(V) and G(V) curves are reported also for K_V_7.2 and K_V_7.3 channels, although no stable intermediate-active state has been observed for these channels. Altogether, K_V_7.4 deviates from K_V_7.2 and K_V_7.3 by displaying a stable intermediate-active voltage sensor state, and deviates from K_V_7.1 by not showing signs of ion conduction at this intermediate state. It would be interesting to investigate whether there are regulatory mechanisms that could convert K_V_7.4 gating, from K_V_7.1/KCNE1-like, to K_V_7.1-like.

Intriguingly, K_V_7.1 channels have a histidine (H240) instead of the “R5”-arginine mutated in the K_V_7.4 R216H variant (Fig.1B). Substitution of H240 in K_V_7.1 to arginine induced a leftward shift in the voltage dependence of the ionic current (40), mirroring the effects of R216H reported here. However, H240R in K_V_7.1 did not apparently change the VSD properties (8). Perhaps the VSD of K_V_7.1 has adapted to be optimally stabilized with a histidine at the “R5” position, such that substitution by arginine does not markedly change its structural dynamics.

The R216 (S4)/E146 (S2) interaction (Fig.5) is observed also in the K_V_7.2 VSD, involving homologous amino-acids R210 (Fig.1B) and E140, and stabilizing the active VSD conformation (18, 41). Thus, the charge/countercharge interaction network appears to be conserved in at least two members of the K_V_7-channel family.

*In vivo*, K_V_7.4 channels are subject to regulatory interactions with protein partners that tune their biophysical properties. These include inhibition by Ca^2+^-bound calmodulin, which interacts with cytosolic C-terminal helices (42), but also the S2-S3 loop in the VSD (43, 44), and potentiation by the G-protein βγ complex (45). We propose that VCF is the ideal tool to investigate such interactions *in cellula*, as in previous studies of Ca^2+^- and Gβγ-dependent VSD modification in BK_Ca_ and Ca_V_2.2 channels, respectively (46, 47).

Finally, while our work here focused on the properties of K_V_7.4 homo-tetramers, K_V_7.4 subunits can form heteromeric channels with other members of the K_V_7 family, like K_V_7.5 in various smooth-muscle tissue, where K_V_7.4 can also co-assemble with the KCNE4 auxiliary subunit. It would be exciting in future studies to determine how such physiologically relevant interactions impact voltage sensor movement and channel opening.

## Materials and Methods

Please see the supplemental information for the full materials and methods. The use of animals, including the performed surgery to isolate oocytes for electrophysiological studies, was reviewed and approved by the regional board of ethics in Linköping, Sweden (Case no. 1941 and 14515). Cut-open oocyte Vaseline gap (48–50) and voltage-clamp fluorometry (22, 23, 50) experiments were performed as previously described (51), with the addition of pulsed fluorescence excitation to minimize photobleaching (Fig.1E-F).

## Acknowledgments

We dedicate this implementation of pulsed excitation for VCF to Ariel L. Escobar. We are grateful to H. Peter Larsson for comments on the manuscript, and members of the Elinder, Liin and Pantazis groups for expert oocyte preparation. The clone for human K_V_7.4 was a kind gift from Dr. Nicole Schmitt at the University of Copenhagen. The molecular dynamics simulations were enabled by resources provided by the National Academic Infrastructure for Supercomputing in Sweden (NAISS), partially funded by the Swedish Research Council through grant agreement no. 2022-06725. Funding: start-up funds from the Linköping University Wallenberg Center for Molecular Medicine / the Knut and Alice Wallenberg Foundation (A.P.); Hjärnfonden (The Swedish Brain Foundation) grant FO2024-0299 (A.P.); Vetenskapsrådet (The Swedish Research Council) grants 2022-00574 (A.P.) and 2021-01885 (S.I.L.). This work was supported by the Italian Ministry for University and Research (MUR), Next Generation EU, Mission 4, Component 2, CUP E63C22002170007 (Project “A multiscale integrated approach to the study of the nervous system in health and disease”, MNESYS) (to MT and FM).

## Author Contributions

Conceptualization, A.P., M.T., S.I.L.,; Methodology, A.P.; Investigation, M.N., D.J.A.F., A.S.K, K.W., F.M.; Formal Analysis: M.N., D.J.A.F., A.S.K., K.W., A.P., F.M.; Writing –Original Draft, M.N., D.J.A.F., A.S.K., S.I.L., A.P.; Writing –Review & Editing, all; Funding Acquisition, M.T., A.P., S.I.L.; Resources: S.I.L., A.P.; Supervision: S.I.L., M.T., A.P.

## Competing Interest Statement

No competing interests to declare.

## Notes

### Competing Interest Statement

The authors have declared no competing interest.

## References

1. C. Kubisch, et al., KCNQ4, a novel potassium channel expressed in sensory outer hair cells, is mutated in dominant deafness. Cell 96, 437–446 (1999).

2. T. Kharkovets, et al., KCNQ4, a K+ channel mutated in a form of dominant deafness, is expressed in the inner ear and the central auditory pathway. Proc. Natl. Acad. Sci. U. S. A. 97, 4333–4338 (2000).

3. F. A. Iannotti, et al., Expression, localization, and pharmacological role of Kv7 potassium channels in skeletal muscle proliferation, differentiation, and survival after myotoxic insults. J. Pharmacol. Exp. Ther. 332, 811–820 (2010).

4. A. Martelli, et al., Vasorelaxation by hydrogen sulphide involves activation of Kv7 potassium channels. Pharmacol. Res. 70, 27–34 (2013).

5. M. V. Soldovieri, F. Miceli, M. Taglialatela, Driving with no brakes: molecular pathophysiology of Kv7 potassium channels. Physiol. Bethesda Md 26, 365–376 (2011).

6. S. I. Börjesson, F. Elinder, Structure, Function, and Modification of the Voltage Sensor in Voltage-Gated Ion Channels. Cell Biochem. Biophys. 52, 149–174 (2008).

7. J. D. Osteen, et al., KCNE1 alters the voltage sensor movements necessary to open the KCNQ1 channel gate. Proc. Natl. Acad. Sci. U. S. A. 107, 22710–22715 (2010).

8. K. J. Ruscic, et al., IKs channels open slowly because KCNE1 accessory subunits slow the movement of S4 voltage sensors in KCNQ1 pore-forming subunits. Proc. Natl. Acad. Sci. U. S. A. 110, E559–566 (2013).

9. R. Y. Kim, S. A. Pless, H. T. Kurata, PIP2 mediates functional coupling and pharmacology of neuronal KCNQ channels. Proc. Natl. Acad. Sci. U. S. A. 114, E9702–E9711 (2017).

10. P. Hou, et al., Inactivation of KCNQ1 potassium channels reveals dynamic coupling between voltage sensing and pore opening. Nat. Commun. 8, 1730 (2017).

11. R. Barro-Soria, Epilepsy-associated mutations in the voltage sensor of KCNQ3 affect voltage dependence of channel opening. J. Gen. Physiol. 151, 247–257 (2018).

12. P. Hou, et al., Two-stage electro–mechanical coupling of a KV channel in voltage-dependent activation. Nat. Commun. 11, 676 (2020).

13. M. A. Edmond, A. Hinojo-Perez, X. Wu, M. E. Perez Rodriguez, R. Barro-Soria, Distinctive mechanisms of epilepsy-causing mutants discovered by measuring S4 movement in KCNQ2 channels. eLife 11, e77030 (2022).

14. M. Pusch, L. Bertorello, F. Conti, Gating and Flickery Block Differentially Affected by Rubidium in Homomeric KCNQ1 and Heteromeric KCNQ1/KCNE1 Potassium Channels. Biophys. J. 78, 211–226 (2000).

15. M. A. Zaydman, et al., Domain–domain interactions determine the gating, permeation, pharmacology, and subunit modulation of the IKs ion channel. eLife 3, e03606 (2014).

16. N.-D. Yang, et al., Electro-mechanical coupling of KCNQ channels is a target of epilepsy-associated mutations and retigabine. Sci. Adv. 8, eabo3625 (2022).

17. S. I. Liin, J. E. Larsson, R. Barro-Soria, B. H. Bentzen, H. P. Larsson, Fatty acid analogue N-arachidonoyl taurine restores function of IKs channels with diverse long QT mutations. eLife 5, e20272 (2016).

18. M. V. Soldovieri, et al., Epileptic Encephalopathy In A Patient With A Novel Variant In The Kv7.2 S2 Transmembrane Segment: Clinical, Genetic, and Functional Features. Int. J. Mol. Sci. 20, 3382 (2019).

19. F. Miceli, M. R. Cilio, M. Taglialatela, F. Bezanilla, Gating currents from neuronal K(V)7.4 channels: general features and correlation with the ionic conductance. Channels Austin Tex 3, 274–283 (2009).

20. K. S. Oh, et al., Overlooked KCNQ4 variants augment the risk of hearing loss. Exp. Mol. Med. 55, 844– 859 (2023).

21. L. M. Mannuzzu, M. M. Moronne, E. Y. Isacoff, Direct physical measure of conformational rearrangement underlying potassium channel gating. Science 271, 213–216 (1996).

22. A. Cha, F. Bezanilla, Characterizing Voltage-Dependent Conformational Changes in the ShakerK+ Channel with Fluorescence. Neuron 19, 1127–1140 (1997).

23. C. S. Gandhi, R. Olcese, The voltage-clamp fluorometry technique. Methods Mol. Biol. Clifton NJ 491, 213–231 (2008).

24. M. Priest, F. Bezanilla, Functional Site-Directed Fluorometry. Adv. Exp. Med. Biol. 869, 55–76 (2015).

25. J. Cowgill, B. Chanda, The contribution of voltage clamp fluorometry to the understanding of channel and transporter mechanisms. J. Gen. Physiol. 151, 1163–1172 (2019).

26. S. Bhat, R. Blunck, Characterising ion channel structure and dynamics using fluorescence spectroscopy techniques. Biochem. Soc. Trans. 50, 1427–1445 (2022).

27. A. Pantazis, K. Westerberg, T. Althoff, J. Abramson, R. Olcese, Harnessing photoinduced electron transfer to optically determine protein sub-nanoscale atomic distances. Nat. Commun. 9, 4738 (2018).

28. T. Li, et al., Structural Basis for the Modulation of Human KCNQ4 by Small-Molecule Drugs. Mol. Cell 81, 25-37.e4 (2021).

29. X. Tao, A. Lee, W. Limapichat, D. A. Dougherty, R. MacKinnon, A gating charge transfer center in voltage sensors. Science 328, 67–73 (2010).

30. V. Barrese, J. B. Stott, I. A. Greenwood, KCNQ-Encoded Potassium Channels as Therapeutic Targets. Annu. Rev. Pharmacol. Toxicol. 58, 625–648 (2018).

31. P. S. Chadha, et al., Contribution of kv7.4/kv7.5 heteromers to intrinsic and calcitonin gene-related peptide-induced cerebral reactivity. Arterioscler. Thromb. Vasc. Biol. 34, 887–893 (2014).

32. L. I. Brueggemann, et al., Differential protein kinase C-dependent modulation of Kv7.4 and Kv7.5 subunits of vascular Kv7 channels. J. Biol. Chem. 289, 2099–2111 (2014).

33. T. Kharkovets, et al., Mice with altered KCNQ4 K+ channels implicate sensory outer hair cells in human progressive deafness. EMBO J. 25, 642–652 (2006).

34. N. Savalli, A. Kondratiev, L. Toro, R. Olcese, Voltage-dependent conformational changes in human Ca(2+)- and voltage-activated K(+) channel, revealed by voltage-clamp fluorometry. Proc. Natl. Acad. Sci. U. S. A. 103, 12619–12624 (2006).

35. E. Gorraitz, B. A. Hirayama, A. Paz, E. M. Wright, D. D. F. Loo, Active site voltage clamp fluorometry of the sodium glucose cotransporter hSGLT1. Proc. Natl. Acad. Sci. 114, E9980–E9988 (2017).

36. M. N. Wojciechowski, et al., Different fluorescent labels report distinct components of spHCN channel voltage sensor movement. J. Gen. Physiol. 156, e202413559 (2024).

37. R. Mejía-Alvarez, et al., Pulsed local-field fluorescence microscopy: a new approach for measuring cellular signals in the beating heart. Pflugers Arch. 445, 747–758 (2003).

38. A. Durner, E. Durner, A. Nicke, Improved ANAP incorporation and VCF analysis reveal details of P2X7 current facilitation and a limited conformational interplay between ATP binding and the intracellular ballast domain. eLife 12, e82479 (2023).

39. R. Barro-Soria, et al., KCNE1 divides the voltage sensor movement in KCNQ1/KCNE1 channels into two steps. Nat. Commun. 5, 3750 (2014).

40. G. Panaghie, G. W. Abbott, The role of S4 charges in voltage-dependent and voltage-independent KCNQ1 potassium channel complexes. J. Gen. Physiol. 129, 121–133 (2007).

41. F. Miceli, et al., Gating Consequences of Charge Neutralization of Arginine Residues in the S4 Segment of Kv7.2, an Epilepsy-Linked K+ Channel Subunit. Biophys. J. 95, 2254–2264 (2008).

42. A. Chang, et al., A Calmodulin C-Lobe Ca2+-Dependent Switch Governs Kv7 Channel Function. Neuron 97, 836-852.e6 (2018).

43. W. Zhuang, Z. Yan, The S2–S3 Loop of Kv7.4 Channels Is Essential for Calmodulin Regulation of Channel Activation. Front. Physiol. 11 (2021).

44. Y. Zheng, et al., Structural insights into the lipid and ligand regulation of a human neuronal KCNQ channel. Neuron 110, 237-247.e4 (2022).

45. J. B. Stott, O. V. Povstyan, G. Carr, V. Barrese, I. A. Greenwood, G-protein βγ subunits are positive regulators of Kv7.4 and native vascular Kv7 channel activity. Proc. Natl. Acad. Sci. 112, 6497–6502 (2015).

46. N. Savalli, A. Pantazis, T. Yusifov, D. Sigg, R. Olcese, The Contribution of RCK Domains to Human BK Channel Allosteric Activation *. J. Biol. Chem. 287, 21741–21750 (2012).

47. M. Nilsson, et al., Voltage-dependent G-protein regulation of CaV2.2 (N-type) channels. Sci. Adv. 10, eadp6665 (2024).

48. M. Taglialatela, L. Toro, E. Stefani, Novel voltage clamp to record small, fast currents from ion channels expressed in Xenopus oocytes. Biophys. J. 61, 78–82 (1992).

49. E. Stefani, F. Bezanilla, Cut-open oocyte voltage-clamp technique. Methods Enzymol. 293, 300–318 (1998).

50. A. Pantazis, R. Olcese, “Cut-Open Oocyte Voltage-Clamp Technique” in Encyclopedia of Biophysics, G. Roberts, A. Watts, Eds. (Springer, 2019), pp. 1–9.

51. K. Wang, et al., A rich conformational palette underlies human CaV2.1-channel availability. [Preprint] (2024). Available at: https://www.biorxiv.org/content/10.1101/2024.09.27.615501v1 [Accessed 6 December 2024].

