## Supplemental information for "Two-step voltage-sensor activation of the human K_V_7.4 channel and effect of a deafness-associated mutation"

### Supporting Information Text

#### Materials & Methods

*Oocyte preparation*

Individual oocytes from *Xenopus laevis* frogs were acquired either through surgical removal followed by enzymatic digestion at Linköping University, or purchased from Ecocyte Bioscience (Dortmund, Germany). The use of animals, including the performed surgery, was reviewed and approved by the regional board of ethics in Linköping, Sweden (Case no. 1941 and 14515). Oocytes at developmental stages V-VI were selected for experiments and injected as following.

*Molecular biology*

Human *KCNQ4* (GenBank accession no. NM_004700) inserted into a pXOOM vector was used in this study. cRNA for injection was prepared from DNA using a T7 mMachine transcription kit (Invitrogen, Stockholm, Sweden), and cRNA concentrations were determined spectrophotometrically (NanoDrop 2000c, Thermo Scientific, Stockholm, Sweden). Mutations were introduced using the primers reported in Table 1, through site-directed mutagenesis (QuikChange II XL, with 10 XL Gold cells; Agilent Technologies, Kista, Sweden) and confirmed by sequencing at the Linköping University Molecular Biology unit of the Core Facility.

*Two-electrode voltage clamp*

Oocytes from developmental stages V-VI were injected with 50 ng of cRNA encoding either wild-type or mutant *KCNQ4* and incubated at 16 °C for 2-4 days prior to electrophysiological experiments. Two-electrode voltage clamp experiments were performed using a Dagan CA-1B amplifier (Dagan, MN, USA). Whole-cell $K^+$ currents were sampled using Clampex software (Molecular Devices, San Jose, CA, USA) at 5 kHz and filtered at 500 Hz. All experiments were carried out at room temperature (approx. 20°C). The extracellular recording solution contained (in mM): 88 NaCl, 1 KCl, 0.4 $CaCl_2$, 0.8 $MgCl_2$, and 15 HEPES. pH was set to 7.4 by addition of NaOH. Voltage-step protocols consisted of a −80 mV holding potential and a hyperpolarizing −100 mV pre-pulse for 2 s to ensure channel closure. This was followed by test pulses ranging from −100 mV to +60 mV with incremental depolarizing steps of 10 mV for 3 seconds, to generate $K^+$ currents. The tail pulse was set to −30 mV for 2 s. Data were recorded under constant superfusion (rate set to 1.0 mL/s) with the extracellular recording solution.

*‘Cut-open oocyte Vaseline gap (COVG)*

We used the cut-open oocyte Vaseline gap (COVG), a low-noise, fast voltage-clamp technique (1–3). Each oocyte received 50 ng of $K_V$7.4 cRNA. Injected oocytes were incubated at 16 °C for 3-5 days prior to electrophysiological experiments. The oocyte was placed in a triple-compartment acrylic chamber, with a diameter of 600 µm for the top and bottom apertures. We used following solutions: external solution (mM): 120 Na-methanesulfonate (MES), 1 K-MES, 2 $Ca(MES)_2$, 10 HEPES (pH=7.0); internal solution (mM): 120 K-glutamate, 10 HEPES (pH=7.0); intracellular micro-pipette solution (mM): 2700 Na-MES, 10 NaCl. Low access resistance to the oocyte interior was obtained by permeabilizing the oocyte with 0.1% saponin carried by the internal solution. In order to evoke responses to measure G(V) relationship, each sweep was 5 s in duration with 10 s intersweep interval. The holding potential was –120 mV. Each sweep was composed of a 500-ms prepulse at the holding potential, followed by a three-second step increasing by 10 mV for each sweep from the holding potential. This was followed by a 0 mV step (200 ms), and finally a 1.5-s step at the holding potential. To measure deactivation kinetics, a prepulse consisting of holding the cell at –120 mV for 500 ms and then to 40 mV for 1 s was applied before every sweep. Acquisition then started and the cell was kept for another 500 ms at 40 mV, followed by a three-second step increasing by 10 mV for each sweep from the holding potential (–120 mV) to 0 mV. Every sweep was followed by 500 ms at the holding potential. The intersweep interval was 10 s. The current signal was filtered at 200 Hz and acquired at 1 kHz.

*Voltage-clamp fluorometry*

Voltage-clamp fluorometry was performed by adding epifluorescence and light detection capability to the COVG system (3–5). Prior to mounting on the COVG apparatus, oocytes were stained on ice for 30 min with 100 µM Alexa Fluor 488 $C_5$ Maleimide fluorophore (Thermo Fischer) or for 5 minutes with 20 µM 6-TAMRA C6 maleimide fluorophore (AAT Bioquest), in a depolarizing solution (in mM: 120 K-MeS, 2 $Ca(MeS)_2$, 10 HEPES, pH 7.0), on ice, in the dark, to label the introduced Cys (N196C) at the S4 extracellular flank. The oocytes were then rinsed in dye-free SOS prior to being mounted in the recording chamber. Fluorescence emission and ionic current were simultaneously measured from the same area of membrane isolated by the top chamber. The same electrophysiological apparatus and solutions were used as above, for COVG. The optical set-up, comprising Thorlabs LED light source, optics and photodiode, Open Ephys LED driver, Semrock light filters, and a FEMTO transimpedance amplifier, was as described previously (6).

We used pulsed excitation of the fluorophore to reduce photobleaching of the fluorescence signal during long-lasting protocols. Data were sampled at 10 kHz. The LED driver was triggered by a 5 V analog signal from the digitizer. In Clampex, the LED trigger analog signal output was set to

“Train” with train period 10 samples and pulse width 1 sample. This resulted in a 1-kHz train rate with pulse width 100 μs: a 10% duty cycle. This produced a 10-KHz record where one in every ten samples contained fluorescence emission, and nine samples were acquired in the dark. After acquisition, data were decimated 10-fold in MATLAB. This resulted in a 1-kHz trace including only light samples. The resulting signal had similar background fluorescence as when emission was on constantly, with a fraction of photobleaching (Fig.1E,F). A record without voltage pulses (–120 mV for the same duration as the experimental protocol) provided a trace with only photobleaching. This could be well fit by an exponential decay function with 2 or 3 terms (Clampfit). The fitted curve was then subtracted from the signal to completely remove residual bleaching.

*Electrophysiological data analysis*

TEVC recordings were analyzed using Clampfit 11 (Molecular Devices), by plotting the immediate tail currents (recorded upon stepping to the tail voltage) against the preceding test voltages and fitting data with a Boltzmann function:

$$I_{\text{tail}}(V) = \frac{I_{\text{tail,max}} - I_{\text{tail,min}}}{1 + \exp\left(\frac{V_{0.5} - V}{k}\right)} + I_{\text{tail,min}} \qquad \text{(eq.1)}$$

where $V$ was the voltage of the test-pulse preceding the tail-pulse; $V_{0.5}$ was the half-activation potential; $I_{\text{tail,max}}$ and $I_{\text{tail,min}}$ were the limiting maximal and minimal tail currents, respectively; and $k$ was the slope.

COVG-VCF recordings were analyzed in MATLAB and Excel. Tail currents were fit to the Boltzmann distribution:

$$I_{\text{tail}}(V) = \frac{I_{\text{tail,max}} - I_{\text{tail,min}}}{1 + \exp\left(\frac{z\mathrm{F}}{R\mathrm{T}}(V_{0.5} - V)\right)} + I_{\text{tail,min}} \qquad \text{(eq.2)}$$

where $z$ was the effective valence, $T$ was the room temperature in K, and F and R the Faraday and Gas constants, respectively.

Bleaching was subtracted from fluorescence data in MATLAB after decimation. The $\Delta F$ was calculated by averaging the last 100 ms of the test pulse and fit to either a single ($i$ = 1) or the sum of two ($i$ = 2) Boltzmann distributions:

$$\Delta F(V) = \sum_{n=1}^{i} \frac{\Delta F_{\text{pos},n} - \Delta F_{\text{neg},n}}{1 + \exp\left(\frac{z_n\mathrm{F}}{R\mathrm{T}}\left(V_{0.5,n} - V\right)\right)} + \Delta F_{\text{neg},n} \qquad \text{(eq.3)}$$

Where $\Delta F_{\text{pos}}$ and $\Delta F_{\text{neg}}$ were the $\Delta F$ asymptotes as $V$ approached plus or minus infinity, respectively.

For the sum of two Boltzmann functions, the following constraints were used while fitting:

a) $\Delta F_{pos,2}$ was constrained to be equal to $\Delta F_{neg,1}$
b) $V_{0.5,2}$ was constrained to be equal to the $V_{0.5}$ of channel opening from the same cell
c) $z_2$ was constrained to be equal to the $z$ of channel opening from the same cell

Fitting was performed by least squares using *Solver* in Microsoft Excel.

Fractional amplitude ($f$) for component $n$ (1 or 2) was calculated as:

$$f_n = \frac{|\Delta F_{\text{neg},n} - \Delta F_{\text{pos},n}|}{|\Delta F_{\text{neg},1} - \Delta F_{\text{pos},1}| + |\Delta F_{\text{neg},2} - \Delta F_{\text{pos},2}|} \quad \text{(eq.4)}$$

Channel opening and closing kinetics were determined by fitting traces to the *Exponential standard* function, with one ($i$ = 1) or the sum of two ($i$ = 2) components in Clampfit 11 (Molecular Devices):

$$f\ (t) = C + \sum_{n=1}^{i} A_n \exp(-t\ /\ \tau_n) \quad \text{(eq.5)}$$

where $A$ was the amplitude of the exponential component, $\tau$ was the time constant, and $C$ was an offset.

$\Delta F$ kinetics were determined in MATLAB using *lsqcurvefit* using eq.5 with $i$ = 2 and the following constraints:

a) $A_1$ was constrained to be equal to $-(C+A_2)$
b) $C$ was constrained to be equal to the $\Delta F$.

Only traces with signal-to-noise ratio above 4 were fit.

*Structure preparation and homology modelling*

The human apo $K_V$7.4 structure (PDB: 7BYL) (7) was prepared by filling in the missing loop between the S3 and S4 helices of the VSD. The missing section between Y367 and D524 corresponds with the cytosolic linker between the HA and HB helices. This section is unresolved in all $K_V$7 structures, yet presents a small spatial gap in the structures; we therefore connected both ends using two glycine residues. An additional model containing the R216H mutation was created. Each $K_V$7.4 subunit was modelled in complex with calmodulin, $Ca^{2+}$ ions were inserted into each calmodulin subunit from the apo state $K_V$7.1 structure (PDB: 6UZZ) (8). Missing loop sections were built using the modeller program (9), 200 models were created in each instance. The models were ranked based on the intersection of the *molpdf* and DOPE scores (10). The top scoring models were then processed using the PROPKA tool to assign tautomeric states to titratable residues at a pH of 7.0 (11). The residue H216 was assigned the +1-charged tautomer where both nitrogen atoms of the imidazole ring are protonated, and this would enable salt bridge interactions with surrounding acidic residues.

*Molecular dynamics simulations*

The CHARMM-GUI webserver (12, 13) was used to generate simulation boxes for each of the models, the box size was set to ~ 140 × 140 × 150 Å. In brief, the proteins were placed in a 1-palmitoyl-2-oleoyl-sn-glycero-3-phosphocholine (POPC) lipid bilayer. Water molecules with 0.15 M KCl were added, including excess $K^+$ ions to neutralise overall charge. All systems were simulated using GROMACS 2021.4 (14) with CHARMM 36M force field parameters (15). All systems were equilibrated using the CHARMM-GUI default equilibration protocol which includes gradually withdrawing positional restraints from protein backbone and sidechain atoms. The protocol was slightly modified such that positional restraints of 100 kJ $mol^{-1}$ $nm^{-1}$ were maintained on protein sidechain and backbone atoms for an additional 5 ns, then restraints were withdrawn from sidechain atoms but maintained on backbone atoms for a final 10 ns. Following equilibration, each system was simulated for 500 ns in triplicate with a 2 fs timestep. The particle mesh Ewald method (16) was used to calculate long range electrostatic interactions, the LINCS algorithm was used to constrain hydrogen-bond lengths (17). The pressure was maintained at 1 bar with semi-isotropic pressure coupling using the Berendsen barostat ($\tau_P$ =1.0 ps and isothermal compressibility = $4.5\times10^{-5}$ bar), and the temperature was maintained at 300 K using the v-rescale thermostat ($\tau_T$ = 0.1 ps) (18).

Preliminary simulation analysis revealed that the inclusion of $PIP_2$ molecules mitigated the impact of R216H mutant on S4 helix movement, this is consistent with PIP2 having a role in stabilizing the activated conformation of the VSD (19). Therefore, we analyzed simulations that did not include $PIP_2$ molecules to better characterize the impact of the R216H mutant on S4 movement. Simulation distance and root mean squared deviation (RMSD) analyses were performed using the MDAnalysis tool (20). Distances were measured between the Cα atoms of F146 and R216 or H216 from their respective systems. RMSD was calculated based on Cα atom positions relative to the initial structure and following superimposition. RMSD was calculated for the transmembrane segments (residues S74-Q332), here, Cα atoms of these residues were superimposed, RMSD was also calculated for the intracellular part of the S4 helix (residues 208-218) in each protein chain, here, the Cα atoms from the S1-S3 helix were superimposed. Molecular interactions involving R216 or H216 were analyzed using the ProLIF tool (21), salt-bridge interactions were calculated based on a 4.5 Å cutoff between the heavy atoms of charged groups in acidic and basic residues. Unless otherwise stated, analyses encompassed simulation frames from 200-500ns. Mean ± SEM interaction values are obtained across the 4 voltage sensors and 3 simulation replicas (n=12). Simulations were visualized using PyMOL (The PyMOL Molecular Graphics System, Version 2.5 Schrödinger, LLC).

*Statistics*

Data are reported as mean ± standard error of the mean (SEM), unless otherwise stated. Student's t-test (unpaired, two-tailed) was used to assess significance, and statistical significance was set when *p*-value was <0.05. To compare biophysical properties of $K_V7.4$ cysteine mutants with $K_V7.4$ WT, one-way ANOVA followed by a Dunnett's post hoc test was used, with $K_V7.4$ WT set as the control group.

### Supporting Figures and Tables
### Figure S1

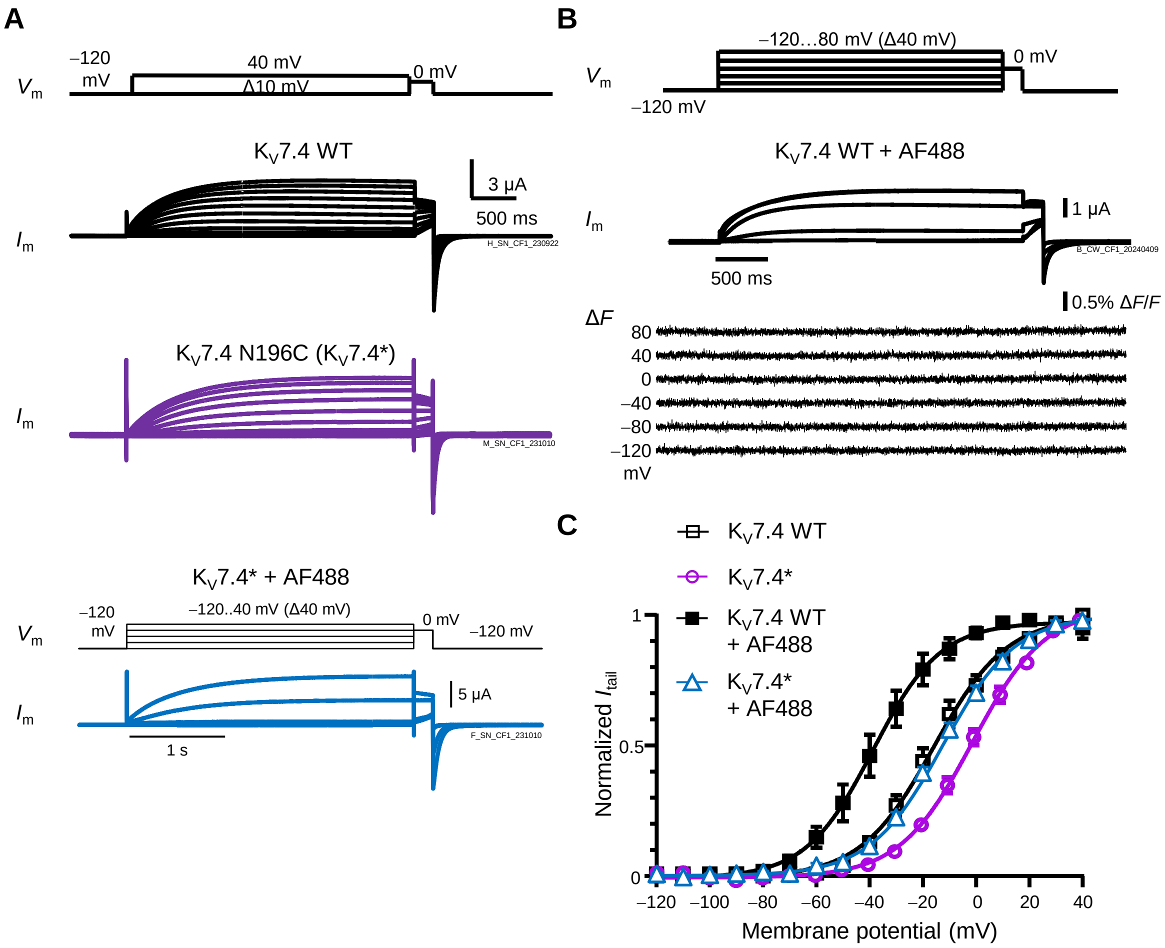


**Figure S1: N196C and AF488 voltage-dependence controls. (A)** Representative current traces from unlabelled $K_V7.4$ WT (black), unlabelled $K_V7.4$ N196C ($K_V7.4$*; purple) and $K_V7.4$* channels labeled with AF488 (blue). The latter is also shown in Fig.2A. **(B)** Representative traces of $K_V7.4$ WT labelled with AF488 showing the voltage protocol ($V_m$) current ($I_m$) and simultaneously acquired fluorescence deflections ($\Delta F$). **(C)** Normalized $I_{tail}(V)$ of unlabelled $K_V7.4$ WT (black open squares): $V_{0.5}$ = –13 ± 2.5; $z$ = 1.9 ± 0.1; $n$ = 8 cells; $K_V7.4$ WT labelled with AF488 (black filled squares): $V_{0.5}$ = –37 ± 3,4; $z$ = 2.3 ± 0.2; $n$ = 6; unlabelled $K_V7.4$* (purple open circles): $V_{0.5}$ = –1.2 ± 2.2; $z$ = 2.0 ± 0.1; $n$ = 12 and $K_V7.4$* labelled with AF488 (blue open triangles): $V_{0.5}$ = –13 ± 1.6 mV; $z$ = 1.9 ± 0.085 $e_0$; $n$ = 11 cells. The latter is also shown in Fig.2C.

### Figure S2

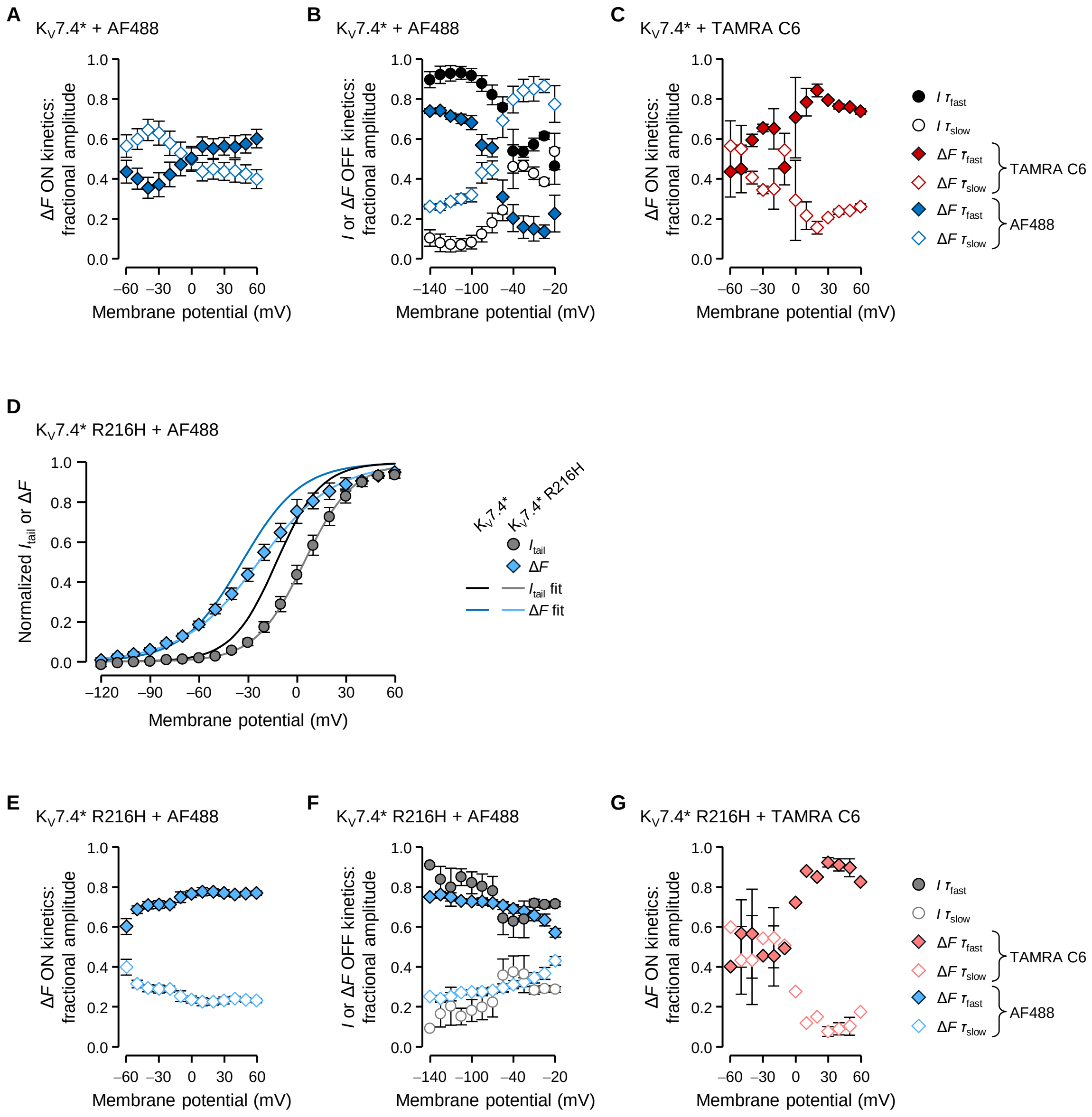


**Figure S2: Supplementary kinetic & voltage-dependence analysis. (A)** Fractional amplitude of kinetic components from activation pulses in $K_V$7.4* channels labeled with AF488 (Fig.2D). **(B)** As in (A), for deactivation pulses (Fig.2E). **(C)** As in (A), for channels labeled with TAMRA C6 (Fig.3C). **(D)** Mean, normalized voltage-dependence of channel opening ($I_{tail}$, filled grey circles) and VSD activation (Δ$F$, filled light-blue diamonds) in $K_V$7.4* R216H channels labelled with AF488 (as in Fig.4E). Curves represent fits to Boltzmann distributions. $I_{tail}$ (grey): $V_{0.5}$ = –4.5 ± 3.2 mV; $z$ = 1.7 ± 0.049 $e_0$; $n$ = 7 cells; Δ$F$ (light blue): $V_{0.5}$ = –24 ± 3.8 mV; $z$ = 1.1 ± 0.099 $e_0$; $n$ = 7. Voltage-dependence curves from $K_V$7.4 channels are shown for reference (Fig.2C): $I_{tail}$ (black); Δ$F$ (blue). **(E-G)** As in (A-C) for channels with R216H (Fig.4G-I).

**Figure S3**

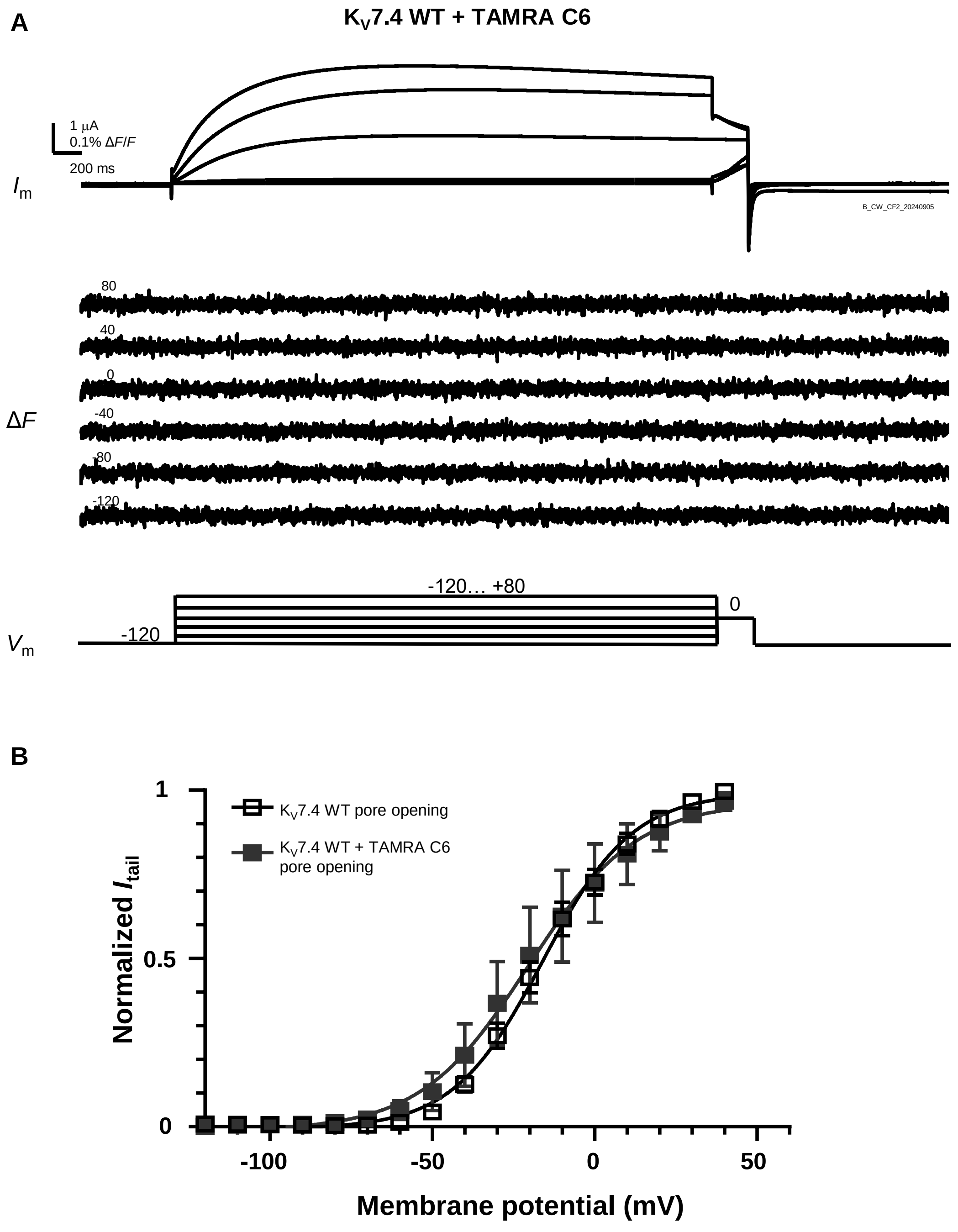


**Figure S3: $K_V$7.4 WT does not report any fluorescence signal when labelled with TAMRA C6.** **(A)** Representative traces of $K_V$7.4 WT labelled with TAMRA C6 showing current ($I_m$), simultaneously acquired fluorescence deflections ($\Delta F$) and membrane potential ($V_m$). **(B)** Normalized voltage dependence of channel opening for $K_V$7.4 WT (black open squares) from Fig.S1C and $K_V$7.4 WT labelled with TAMRA C6 (black filled squares): $V_{0.5} = -9.8 \pm 11$; $z = 1.1 \pm 0.3$; $n = 5$ cells.

**Figure S4**

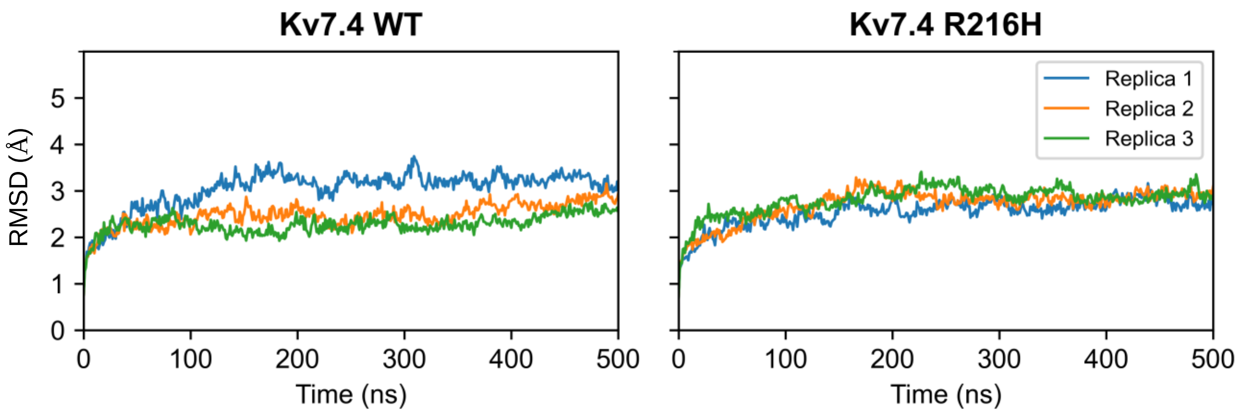


**Figure S4: RMSD of the apo K$_V$7.4 structure with and without the R216H mutation.** RMSD was calculated for the Cα atoms of transmembrane residues 74-332 every 1 ns, following superimposition on the initiation conformation. The average RMSD across the 3 simulations replicas after 200 ns is: 2.7±0.5 Å (WT) and 2.8±0.1 Å (R216H).

**Table S1: Biophysical properties of $K_V$7.4 WT and mutants, as determined by TEVC experiments.**

| **$K_V$7.4 clone** | **$V_{0.5}$ (mV)** | **Adj. *P*** | ***k* (mV)** | **Adj. *P*** | ***n*** |
|---|---|---|---|---|---|
| WT | −9.7 ± 1.9 | - | 12.2 ± 0.4 | - | 6 |
| Q194C | −16.0 ± 1.6 | *0.4380* | 18.2 ± 0.4 | **0.0019** | 8 |
| G195C | 0.4 ± 6.1 | *0.1139* | 26.4 ± 2.9 | **<0.0001** | 6 |
| N196C | −4.0 ± 1.3 | *0.4064* | 11.1 ± 0.3 | *0.8722* | 18 |
| I197C | −38.5 ± 5.2 | **<0.0001** | 20.7 ± 0.9 | **<0.0001** | 5 |
| F198C | −65.2 ± 0.8 | **<0.0001** | 19.7 ± 1.6 | **0.0033** | 3 |

**SI References**


1. M. Taglialatela, L. Toro, E. Stefani, Novel voltage clamp to record small, fast currents from ion channels expressed in Xenopus oocytes. *Biophys. J.* **61**, 78–82 (1992).
2. E. Stefani, F. Bezanilla, Cut-open oocyte voltage-clamp technique. *Methods Enzymol.* **293**, 300–318 (1998).
3. A. Pantazis, R. Olcese, "Cut-Open Oocyte Voltage-Clamp Technique" in *Encyclopedia of Biophysics*, G. Roberts, A. Watts, Eds. (Springer, 2019), pp. 1–9.
4. A. Cha, F. Bezanilla, Characterizing Voltage-Dependent Conformational Changes in the ShakerK+ Channel with Fluorescence. *Neuron* **19**, 1127–1140 (1997).
5. C. S. Gandhi, R. Olcese, The voltage-clamp fluorometry technique. *Methods Mol. Biol. Clifton NJ* **491**, 213–231 (2008).
6. K. Wang, *et al.*, A rich conformational palette underlies human CaV2.1-channel availability. [Preprint] (2024). Available at: https://www.biorxiv.org/content/10.1101/2024.09.27.615501v1 [Accessed 6 December 2024].
7. T. Li, *et al.*, Structural Basis for the Modulation of Human KCNQ4 by Small-Molecule Drugs. *Mol. Cell* **81**, 25-37.e4 (2021).
8. J. Sun, R. MacKinnon, Structural Basis of Human KCNQ1 Modulation and Gating. *Cell* **180**, 340-347.e9 (2020).
9. A. Sali, T. L. Blundell, Comparative protein modelling by satisfaction of spatial restraints. *J. Mol. Biol.* **234**, 779–815 (1993).
10. M.-Y. Shen, A. Sali, Statistical potential for assessment and prediction of protein structures. *Protein Sci. Publ. Protein Soc.* **15**, 2507–2524 (2006).
11. M. H. M. Olsson, C. R. Søndergaard, M. Rostkowski, J. H. Jensen, PROPKA3: Consistent Treatment of Internal and Surface Residues in Empirical pKa Predictions. *J. Chem. Theory Comput.* **7**, 525–537 (2011).
12. S. Jo, T. Kim, V. G. Iyer, W. Im, CHARMM-GUI: a web-based graphical user interface for CHARMM. *J. Comput. Chem.* **29**, 1859–1865 (2008).
13. J. Lee, *et al.*, CHARMM-GUI Input Generator for NAMD, GROMACS, AMBER, OpenMM, and CHARMM/OpenMM Simulations Using the CHARMM36 Additive Force Field. *J. Chem. Theory Comput.* **12**, 405–413 (2016).
14. M. J. Abraham, *et al.*, GROMACS: High performance molecular simulations through multi-level parallelism from laptops to supercomputers. *SoftwareX* **1–2**, 19–25 (2015).
15. J. Huang, *et al.*, CHARMM36m: an improved force field for folded and intrinsically disordered proteins. *Nat. Methods* **14**, 71–73 (2017).
16. U. Essmann, *et al.*, A smooth particle mesh Ewald method. *J. Chem. Phys.* **103**, 8577–8593 (1995).
17. B. Hess, H. Bekker, H. J. C. Berendsen, J. G. E. M. Fraaije, LINCS: A linear constraint solver for molecular simulations. *J. Comput. Chem.* **18**, 1463–1472 (1997).
18. G. Bussi, D. Donadio, M. Parrinello, Canonical sampling through velocity rescaling. *J. Chem. Phys.* **126**, 014101 (2007).
19. V. S. Mandala, R. MacKinnon, The membrane electric field regulates the PIP2-binding site to gate the KCNQ1 channel. *Proc. Natl. Acad. Sci.* **120**, e2301985120 (2023).
20. MDAnalysis: A toolkit for the analysis of molecular dynamics simulations - Michaud-Agrawal - 2011 - Journal of Computational Chemistry - Wiley Online Library. Available at: https://onlinelibrary.wiley.com/doi/full/10.1002/jcc.21787 [Accessed 6 December 2024].
21. C. Bouysset, S. Fiorucci, ProLIF: a library to encode molecular interactions as fingerprints. *J. Cheminformatics* **13**, 72 (2021).
